# GRaphical footprint based Alignment-Free method (GRAFree) for reconstructing evolutionary Traits in Large-Scale Genomic Features

**DOI:** 10.1101/389403

**Authors:** Aritra Mahapatra, Jayanta Mukherjee

## Abstract

In our study, we attempt to extract novel features from mitochondrial genomic sequences reflecting their evolutionary traits by our proposed method GRAFree (GRaphical footprint based Alignment-Free method). These features are used to build a phylogenetic tree given a set of species from insect, fish, bird, and mammal. A novel distance measure in the feature space is proposed for the purpose of reflecting the proximity of these species in the evolutionary processes. The distance function is found to be a metric. We have proposed a three step technique to select a feature vector from the feature space. We have carried out variations of these selected feature vectors for generating multiple hypothesis of these trees and finally we used a consensus based tree merging algorithm to obtain the phylogeny. Experimentations were carried out with 157 species covering four different classes such as, Insecta, Actinopterygii, Aves, and Mammalia. We also introduce a measure of quality of the inferred tree especially when the reference tree is not present. The performance of the output tree can be measured at each clade by considering the presence of each species at the corresponding clade. GRAFree can be applied on any graphical representation of genome to reconstruct the phylogenetic tree. We apply our proposed distance function on the selected feature vectors for three naive methods of graphical representation of genome. The inferred tree reflects some accepted evolutionary traits with a high bootstrap support. This concludes that our proposed distance function can be applied to capture the evolutionary relationships of a large number of both close and distance species using graphical methods.

## Introduction

In studying phylogeny of different species using molecular data, mostly mutations, insertion, and deletion of residues in various homologous segments of DNA sequences are observed by computational biologists[18], [27]. This approach is sensitive to the selection of segments (e.g. genes, coding segments, etc.) of the sequence. Moreover, the homologous segments are very small portions (< 2%) of the whole genome [51]. The roles of majority of the genome sequences (≈ 98%) are unknown. Hence those parts are considered as “*junk*” [15], [49].

The mitochondrial genomes (mtDNA) are relatively simpler than the whole genome. It consists of a limited number of genes, tRNAs, etc. Moreover, the “junk” segments are negligible with respect to the length of the mtDNA (generally ≈ 1%). Most importantly, mtDNA are haploid, inherited maternally in most animals [12], and recombination is very rare event in it [16]. So the changes of mtDNA sequence occur mainly due to mutations.

There are various challenges in using mtDNA sequences in computation and analysis. During the evolution process the genes of mtDNA very often change their order within the mtDNA and also get fragmented [33], [22], [5], [2]. This violates the collinearity of homologous regions very often [77]. The length of the mtDNA as well as the length of genes are also different for different species which makes it difficult to align the homologous regions. Apart from these facts, the complexity, versatility, and the huge length of the data make it difficult to develop any simple method in comparative genomics [46]. Conventional methods compute the distance between sequences through computationally intensive process of multiple sequence alignment [29], which remains a bottleneck in using whole genomic sequences for constructing phylogeny [25]. As a result, there exist a few works which attempt to discover evolutionary features in the larger apparent non-homologous regions of the genomic sequences using alignment-free methods.

The existing alignment-free methods can be broadly categorized into four types:

1. **k-mer/word frequency based methods:** The comparison between two sequences are derived by the variation of the frequency of optimized k-mer. Feature frequency profile (FFP) [62], [63], composition vector (CV) [69], return time distribution (RTD) [30], frequency chaos game representation (FCGR) [28] [23]) used this method.
2. **Substring based methods:** The pairwise distances are measured by the average length of maximum common substrings of two sequences, e.g., average common substring (ACS) [67], average common substring with k-mismatches (ACS-k) [35], mutation distance [24].
3. **Information theory based methods:** The alignment-free sequence comparison method become effective by inheritting different concepts from information theory like entropy, mutual information, etc. Base base correlation [43], Information correlation and partial information correlation (IC-PIC) [20], and Lempel-Ziv compress [50] proposed various information theory based methods to compare two sequences.
4. **Graphical representation based methods:** Here the DNA/amino acid sequences are represented in multidimensional space. The pairwise distances are obtained by comparing the graphs. Iterated map (IM) [1] adopted this technique to compute the distance between two sequences.

Most of the methods have various limitations. They are not suitable to deal with a large number of taxa, and the size of the input sequences is also limited [6, 45, 7]. It is found that an online tool named Alfree [77] can accept the total length of all sequences up to two lakhs. Similarly another online tool CVTree3 [55], [78] works on coding sections only. The offline version of CVTree [55], [78], 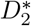 [59], [68] is a very expensive process with respect to both memory and time. More over, the genomic data often works better or increase support for smaller datasets. For the larger dataset of very diverge species the phylogenetic tree construction methods have often failed [54].

Due to these difficulties, conventional methods of phylogenetic reconstruction are restricted to working with whole genome sequences as well as large dataset. For the last three decades, several methods have been introduced to represent the DNA sequence mathematically (both numerically and graphically) [48]. It has been hypothesized that each species carries unique patterns over their DNA sequence which makes a species different from others [34]. Exploration of those distributions for unique characterization is the key motivation behind the mathematical representation of a genome. There exist various representations of the large genome sequences through line graph by mapping the nucleotides to various numeric representations. Considering the genome sequences as the signal (called genomic signal), these methods analyze respective sequences using different signal processing techniques. Several techniques have been proposed to represent DNA sequences graphically in 2D space. One of the way to represent is by considering the structural groups of DNA sequences, such as purine (*A*, *G*) and pyrimidine (*C*, *T*) [47], amino (*A*, *C*) and keto (*T*, *G*) [37], and strong H-bond (C, G) and weak H-bond (*A*, *T*) [21]. The graphical representation has inherent a serious limitation of overlapping paths which causes loss of information [56]. In some techniques, the sequence is represented as an entity in higher dimensions such as, in 3-D [57, 39, 9, 26, 42], 4-D [11], 5-D [40], and 6-D [41]. The increase of dimension reduces the probability of occurrence of degeneracy, but it causes difficulty in visualization. In few schemes, like Worm Curve [58], DV-Curve [75], cell representation [71], etc., the DNA sequences are represented in a non-overlapping fashion.

GRAFree can be applied on any graphical method. GRAFree also lifts the loss of information due to overlapping paths by considering the coordinates of each nucleotide. In this study, we consider three sets of structural groups of nucleotides (purine, pyrimidine), (amino, keto), and (weak H-bond, strong H-bond) separately for representing DNA by a sequence of points in a 2-D integral coordinate space. This point set is called **Graphical Foot Print (GFP)** of a DNA sequence. We propose a technique for extracting features from GFPs and use them for constructing phylogenetic trees. As there are three different types of numerical representation of nucleotides, there are three different hypotheses for the phylogeny. Each of them is found to be statistically significant compared to a tree randomly generated. We also generate a consensus tree from these three hypotheses by applying a tree merging algorithm called COSPEDTree [3, 4].

Experimentations were carried out with a large dataset of total 157 species from four different classes, namely, Insecta (insect), Actinopterygii (ray-finned fish), Aves (bird), and Mammalia (Mammal).

The contributions made in this work are highlighted below:

- Revisiting the concept of Graphical Foot Print (GFP), 2D representation of DNA sequences, and introducing the concept of drift in GFP, which is found to be translation invariant for a sequence.
- Representation of a fragment of drift by a novel 5 dimensional descriptor. All the 5-D descriptors together represent a genotype characteristic for a species.
- Proposed a new distance function to measure the dissimilarity among species, and use the distance matrix for generating phyolgenetic tree by distance based methods such as UPGMA [64]. The distance function is proved to be a metric.
- Proposed a technique to select the value of the parameters involved in computing the distance matrix.

## 1 Materials and Methods

### Feature space

#### Definition 1. Graphical Foot Print (GFP)

*Let a sequence*, 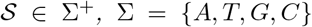. *For each combination of Purine (R)/Pyrimidine (Y), Amino (M)/Keto (K), and Strong H-bond (S)/Weak H-bond (W),the GFP of* 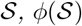, *is the locus of 2-D points in an integral coordinate space, such that (x_i_,y_i_) is the coordinate of the alphabet* 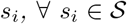 *for i* = 1, 2, …, *n*, *and x*_0_ = *y*_0_ = 0.

*Case-1: for Purine/Pyrimidine*

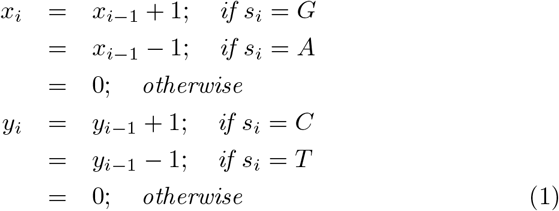
*Case-2: for Strong H-bond/Weak H-bond*

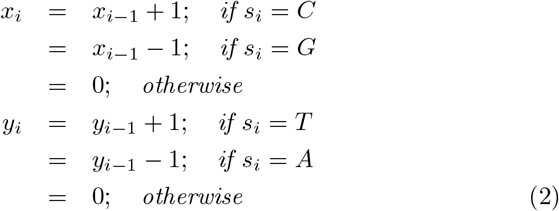
*Case-3: for Amino/Keto*

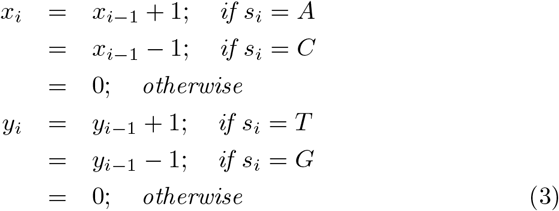

We denote GFPs of Case-1, Case-2 and Case-3, as GFP-RY (Φ_*RY*_), GFP-SW (Φ_*SW*_) and GFP-MK (Φ_*MK*_), respectively.

#### Definition 2. Drift of GFP

*Let* 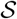 *be a DNA sequence and s_i_ be the alphabet (s_i_* ∈ {*A*,*T*,*G*,*C*}*) at the i^th^ position of* 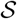. *Let* 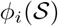 *denote the corresponding (x_i_,y_i_) the coordinate of s_i_ in* 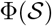.

*Then for length *L*, drift at the i^th^ position is defined as*, 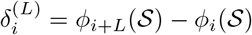, where 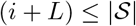

*Considering the drifts for every i^th^ location of the whole sequence, the sequence of drifts is denoted by*

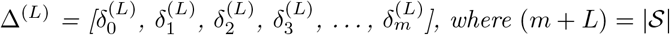

For GFP-RY (refer to Definition 1), an element (*x_i_*, *y_i_*) in 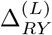 provides excess numbers of *G* from *A* and *C* from *T* in segment of length *L* starting from the *i*^th^ location, respectively. Similarly, in GFP-SW, they are the excess numbers of *C* from *G* and *T* from *A* (represents as 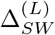), and in GFP-MK, they correspond to the excess numbers of *A* from *C* and *T* from *G* (represents as 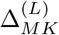), respectively.

We also call the elements of Δ^(*L*)^ as points, as they can be plotted on a 2-D coordinate system. We call this plot as the scatter plot of the drift sequence. Similarly, we get a scatter plot of a GFP. Compared to 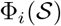, Δ^(*L*)^ is translation invariant as its set of points does not depend on the starting point of the sequence. It has been observed that in many cases the scatter plots of Δ have similar structure for closely spaced species mentioned in literature. In Fig. 1 we demonstrate the scatter plots of GFPs and drift sequences of two species from each class namely, *Drepanotermes* sp. and *Macrognathotermes errator* from insects, *Bathygadus antrodes* and *Bregmaceros nectabanus* from fishes, *Jacana jacana* and *Raphus cucullatus* from birds, *Canis familiaris* and *Panthera tigris tigris* from mammals. It can be observed that the species from same class (insect, fish, bird, or mammal) have the similar pattern in their drift sequences which intuitively indicates that the intraclass species are closer that the interclass species. It can also be observed that differences between two GFPs get reflected in their respective drifts. It is noted that the GFPs of *Bathygadus antrodes, Bregmaceros nectabanus*, and *Canis familiaris* (refer to Fig. 1 (c, d, g), respectively) have the similar patters, where as their drifts, shown in Fig. 1 (k, l, o), respectively, are quite different.

**Figure 1:**
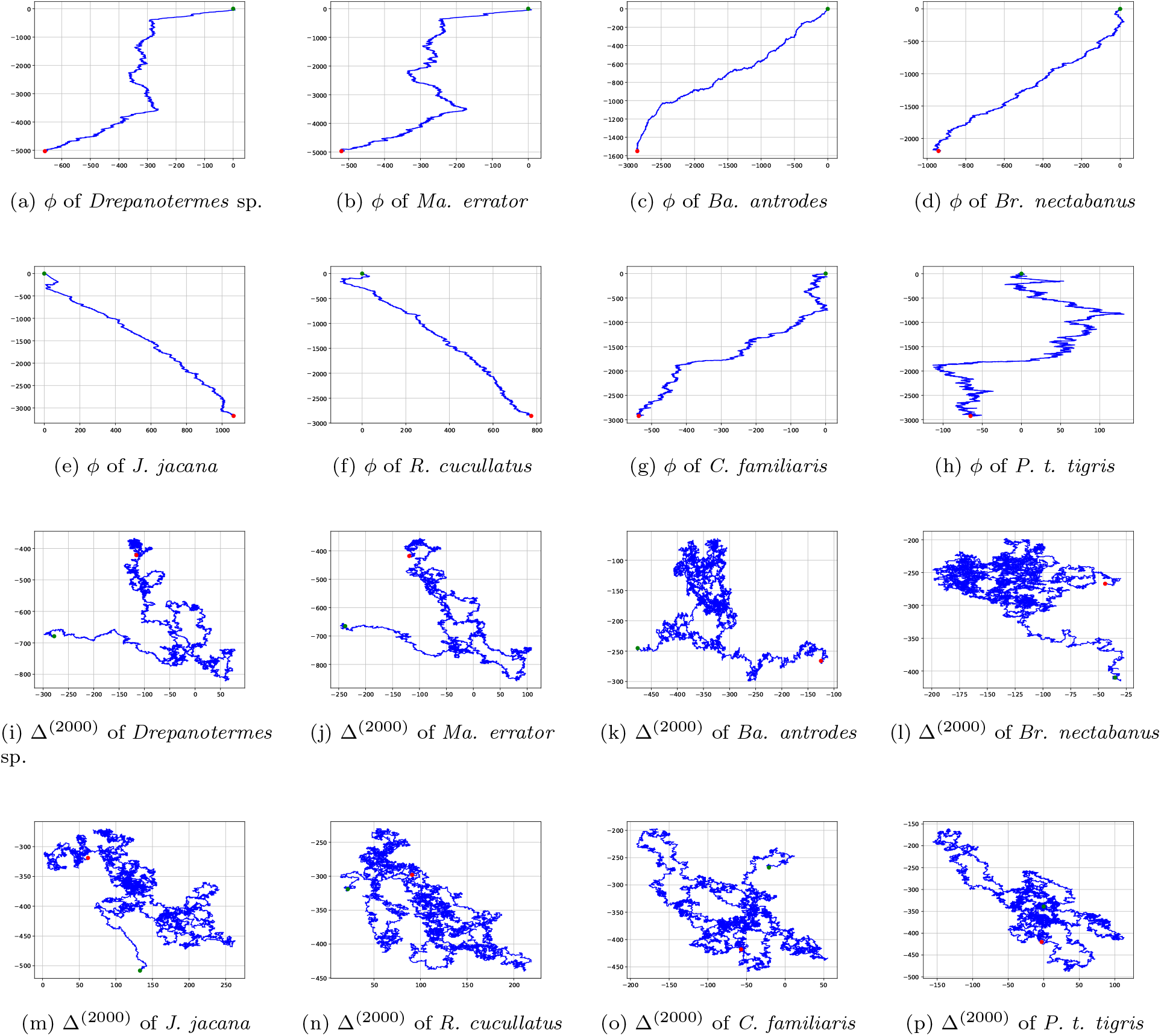
Φ_*RY*_ and Δ_*RY*_ of few species from our dataset. Fig. 1 (a, b) are the Φ_*RY*_ generated from the insects namely, *Drepanotermes* sp. and *Macrognathotermes errator*, respectively. Fig. 1 (c, d) are the Φ_*RY*_ generated from the fishes namely, *Bathygadus antrodes* and *Bregmaceros nectabanus*, respectively. Fig. 1 (e, f) are the *Φry* generated from the birds namely, *Jacana jacana* and *Raphus cucullatus*, respectively. Fig. 1 (g, h) are the Φ_*RY*_ generated from the mammals namely, *Canis familiaris* and *Panthera tigris tigris*, respectively. Fig. 1 (i-p) are the Δ_*RY*_ for *L* − 2000 of the corresponding species. The green and red dots are the start and end points of the graph, respectively.

We represent spatial distribution of these points of Δ by an elliptical model using a five dimensional feature descriptor: (*μ*, Λ, λ, *θ*), where *μ* = (*μ_x_,μ_y_*) is the center of the coordinates, Λ and λ are major and minor eigen values of the covariance matrix, and *θ* is the angle formed by the eigen vector corresponding to Λ with respect to the *x*-axis. We make 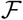 number of non overlapping equal length fragments from Δ and represent each fragment using the five dimensional feature descriptor.

### Distance function and its properties

For two sequences 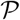 and 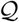 with the feature descriptors of *i^th^* fragments 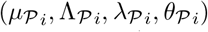 and 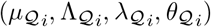, where 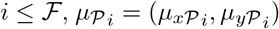 and 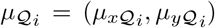, we propose the following distance function between them,

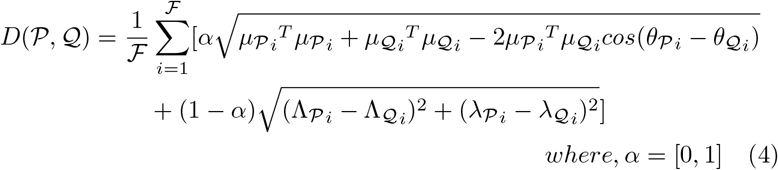

#### Lemma 1.

*The distance D between two sequences is a metric.*

*For any three sequences, 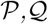 and 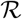, we have*

1. *Non-negativity*. 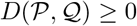
2. *Identity.* 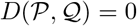 *if and only if* 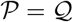
3. *Symmetry.* 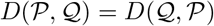
4. *Triangle inequality.* 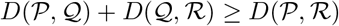

*Proof.* The properties 1, 2 and 3 can be proved from the definition itself. Here we prove the property 4.

The distance between a single fragment of 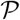 and 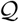 is,

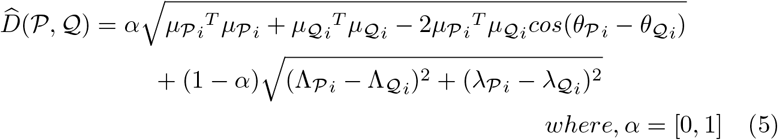

The distance between a single fragment of 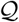 and 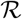 is,

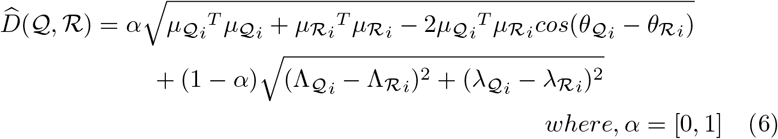

The distance between a single fragment of 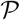 and 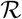 is,

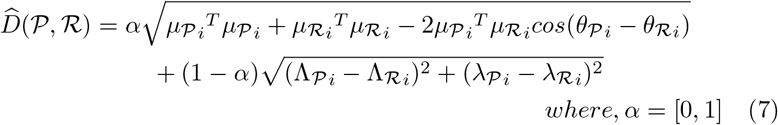

Let,

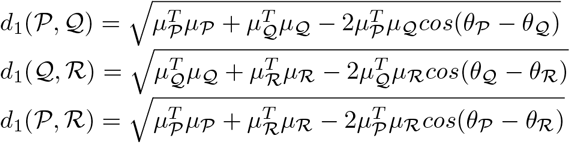

First, we prove,

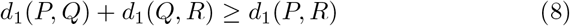

Fig. 2 represents the idea behind computing the distance between two sequences. So, from the figure we observe that 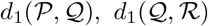 and 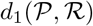 form a triangle. Using triangulation inequality Eq. (8) can be proved.

For single fragment Let,

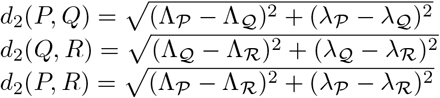

Similarly, it can be proved that,

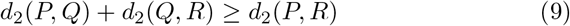

Hence, by combining Eq. (8) and Eq. (9) it can be said that, 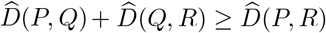.

Hence, 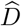 is a metric. As *D* is the linear combination of 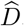. So *D* is also be a metric.

**Figure 2:**
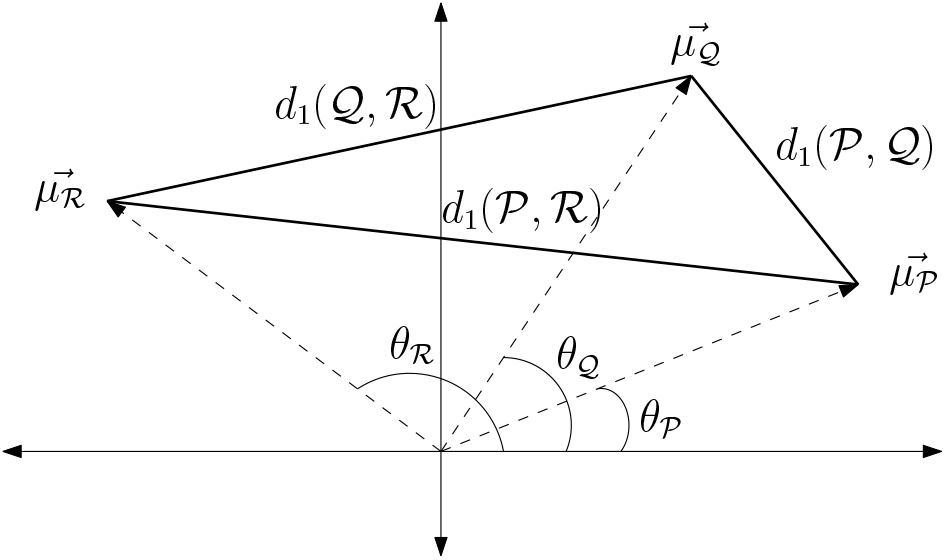
Computation of 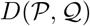

### Taxon sampling and acquiring mitochondrial genome

We have selected various mitochondrial genome sequences sequenced by various researchers such as insects are selected from [8], [65], ray-finned fishes are selected from [61], [76], Aves data are selected from [19], [32], [53], and Mammalian data are selected from [31], [73], [74] [52]. We ignore those accession numbers which store some selected genes of mtDNA. Hence, we have studied over 157 species of four different classes - Insecta (insect), Actinopterygii (ray-finned fish), Aves (bird), and Mammalia (Mammal). The selected data have been downloaded from the NCBI database^1^. The average percentage of unrecognized nucleotide of all 157 mtDNA is 0.06% which inferred that the data we selected for this study are quite good in quality. Details of all the species are listed in Table 1.

**Table 1:**
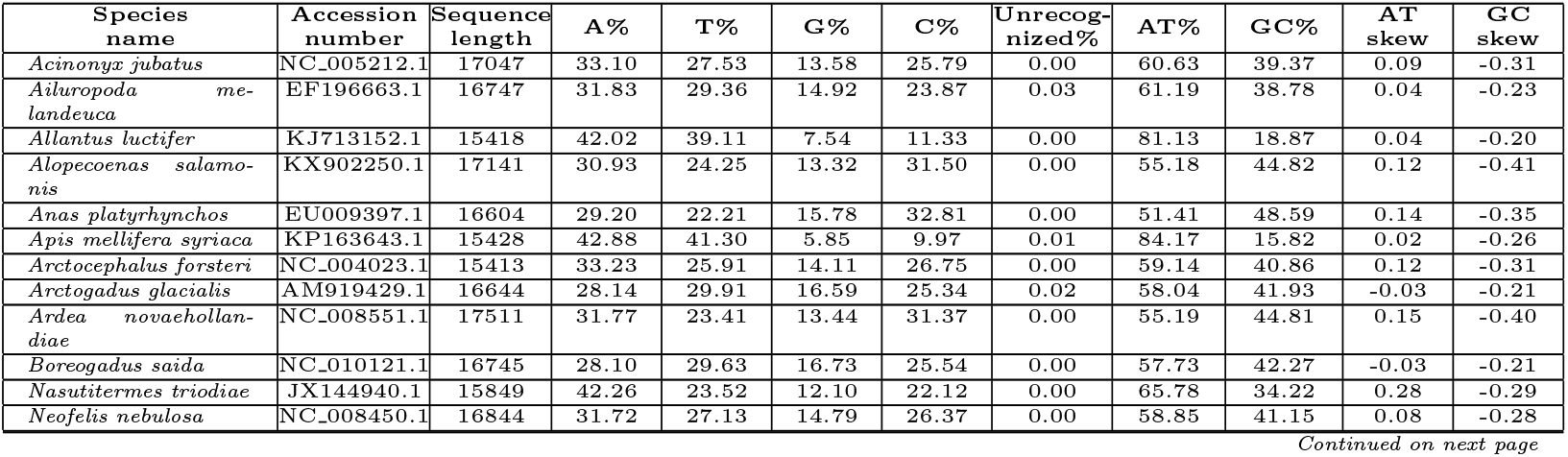
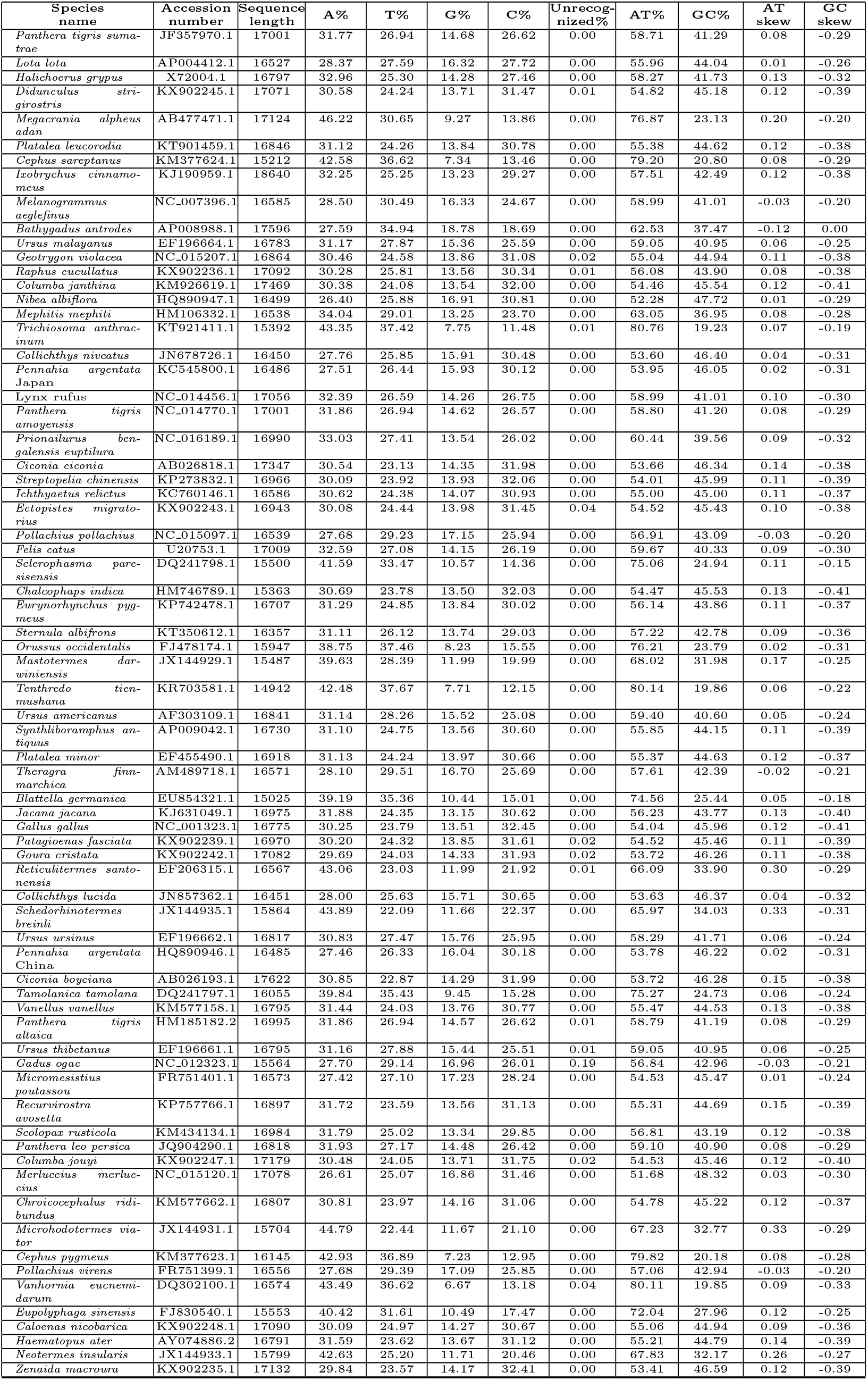
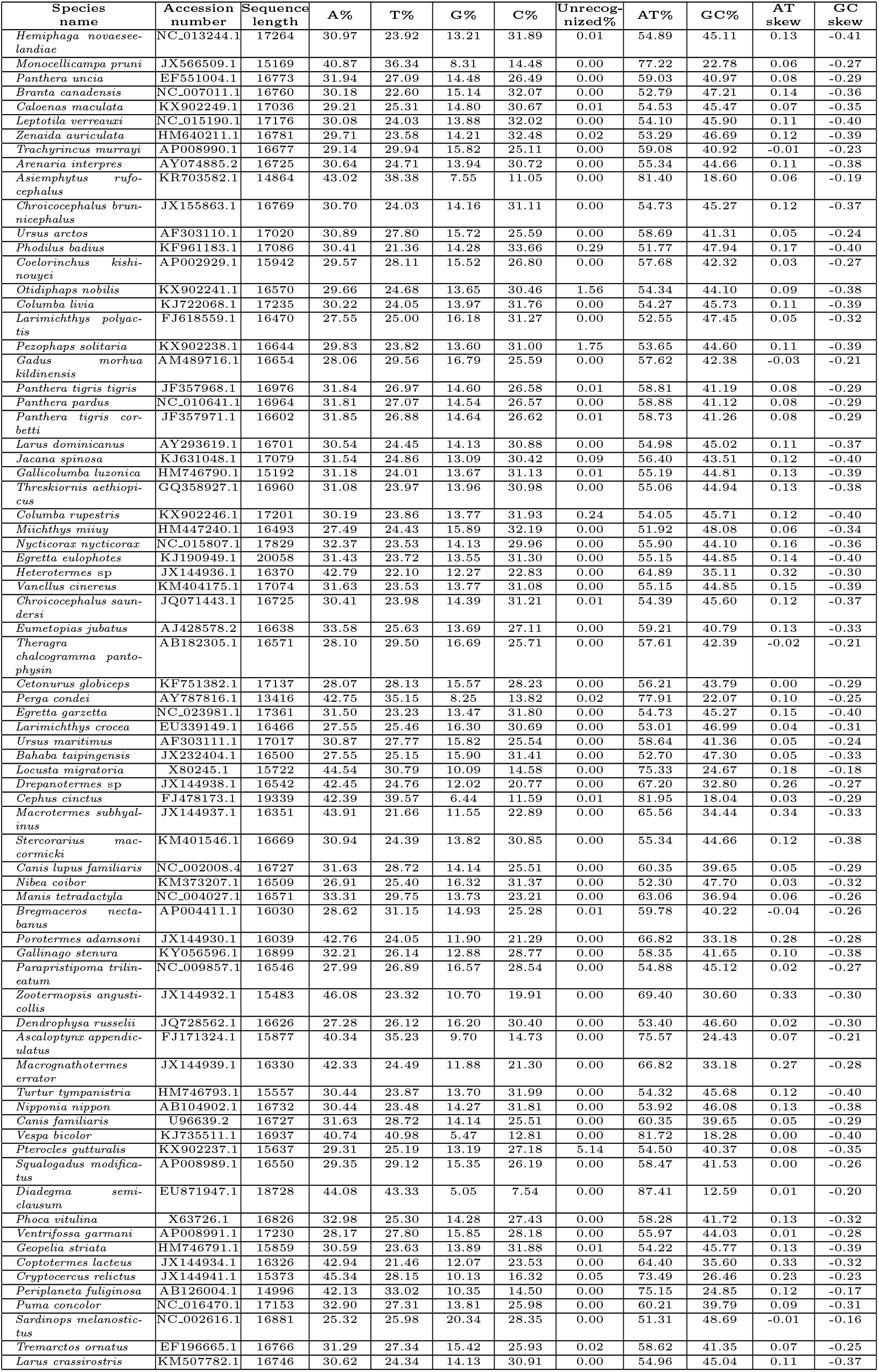
List of species

### Data acquisition

We have developed a python based web scraper which can download the whole genome sequences, gene, and amino acid sequences (from the NCBI database server) by specifying either the accession numbers of the target species, or the name of a particular gene. In the former case, the whole genome sequences, different gene and amino acid sequences of the query species are downloaded. In the second case, all of the homologs of the query gene are extracted from the server. The web scraper extracts information from the NCBI data repository using *Entrez Global Query Cross-Database Search System*^2^. Entrez is a primary text search and retrieval system of NCBI database. The search system provides nine e-utilities, out of which “ESearch”, “ELink”, and “EFetch” have been used in our tool. From the downloaded items we consider only the mitochondrial genome sequences used in the subsequent analysis. Accession numbers of the individual species used in the current study are listed in Table 1.

### Phylogenetic inference

We apply the proposed distance measure (refer to Eq. (4)) over 157 selected species from four classes to compute pairwise distances between them with different values of length, *L* (from 50 to 5000), 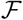 (from 1 to 200), and *α* [0,1]. We compute all the feature sets separately for GFP-RY, GFP-SW, and GFP-MK (refer to Eq. (1), (2), (3), respectively). By applying Unweighted Pair Group Method with Arithmetic Mean (UPGMA) [64] over these distance matrices, we get the phylogenetic tree for each such case. The inferred phylogenetic trees for GFP-RY, GFP-SW, and GFP-MK are represented as 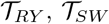, and 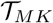, respectively.

Finally, given 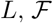, and *α*, we get three phylogenetic trees for GFP-RY, GFP-SW, and GFP-MK, 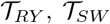, and 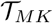, respectively. These trees are combined following a consensus tree merging algorithm (COSPEDTree-II [4]) and get 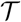.

### Selecting parameter values

We apply following three step technique to select the 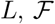, and *α*.

1. Selecting the value of *L* using Shannon entropy of the sequence of each species.
2. Considering the intraclass variances and interclass distances of the features of each species to select the value of 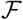.
3. By considering the same for the pairwise distances we select the value of *α*.

It is empirically noticed that the selection criteria we proposed derive the trees 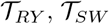, and 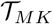 infer better clades with the four different classes of species.

### Selection of the value of *L*

Shannon entropy [60] is used to measure the randomness in genomic data [66]. For different values of *L* (from 50 to 5000 with the difference of 50), we compute the Shannon entropy of the drift sequence (Δ) of individual species. The high value of entropy infers that for the value of *L*, the Δ^(*L*)^ contains high number of unique point coordinates. Fig. 3 shows that initially the entropy of all the species increase by increasing *L*. At almost *L* = 800, the entropy of all the species become stabilized at a high level. By increasing the value of *L* after that does not change the entropy at any significant level. From Fig. 3 it can be noted that for *L* ≥ 800, the Δ^(*L*)^ contains significantly large number of unique point coordinates than that of *L* < 1800. It is also be pointed out that increasing the value of *L* reduces the number of point coordinates in the corresponding Δ. Hence, we choose the value of *L* as 800.

**Figure 3:**
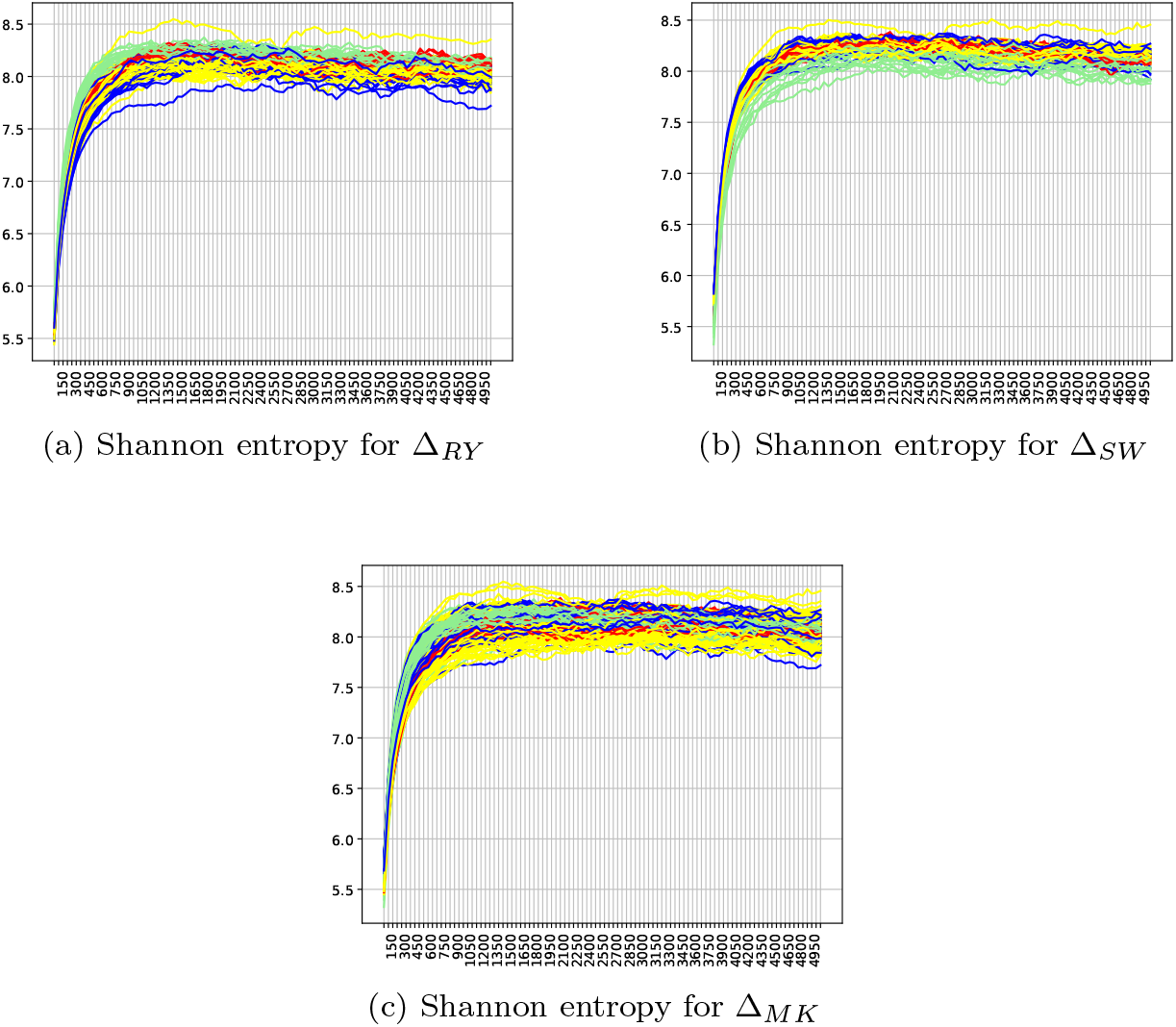
Shannon entropy for all the value of *L* from 50 to 5000. Different color graphs represent the Shannon entropy of four different types of species, Mammalia (Red), Aves (Yellow), Actinopterygii (Blue), and Insecta (Green).

### Selection of the value of 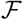

Here we consider the feature vector for each species, say *χ_s_*, where s is a species. By applying the distance metric (refer to Eq. (4)), we compute the distances, say, *D*(*χ_si_*,*μ_i_*), where *χ_si_* is the feature vector of the species s selected from class *i*, and μ_i_ is the mean of the feature vector of all species from class *i*. The variance of the computed distances of class *i*, say, 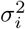, represents the separation of the feature vectors of intraclass species.

So, for *C* number of classes the mean of intraclass variances,

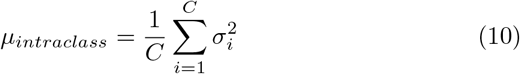

To derive the interclass distances, we consider the *μ*, where,

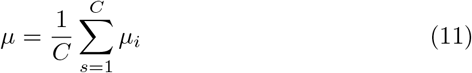

By applying the distance metric (refer to Eq. (4)) we compute *D*(*μ_i_*, *μ*) which represents the separation of the feature vectors of interclass species.

So, for *C* number of classes the mean of interclass distance,

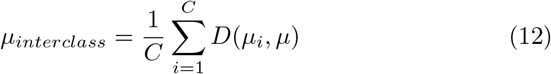

Using this two elements we derive the discriminant score of the selected species as,

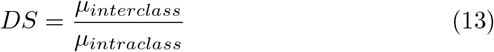

Maximizing the *DS* is equivalent to getting a good separation between the feature vectors of interclass species. We apply this method for different values of 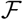 (from 1 to 200) and the optimized values of *L*, here it is *L* ≥ 800. It is found from Fig. 4 (a, c, and e) that for all the values of *L*, the value of *DS* increases with an increasing value of 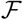. It can be noticed that the overall value of *DS* is maximum for *L* = 800. So to select the value of 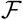, we consider 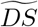 for *L* = 800, where,

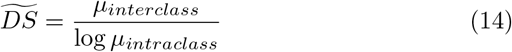

**Figure 4:**
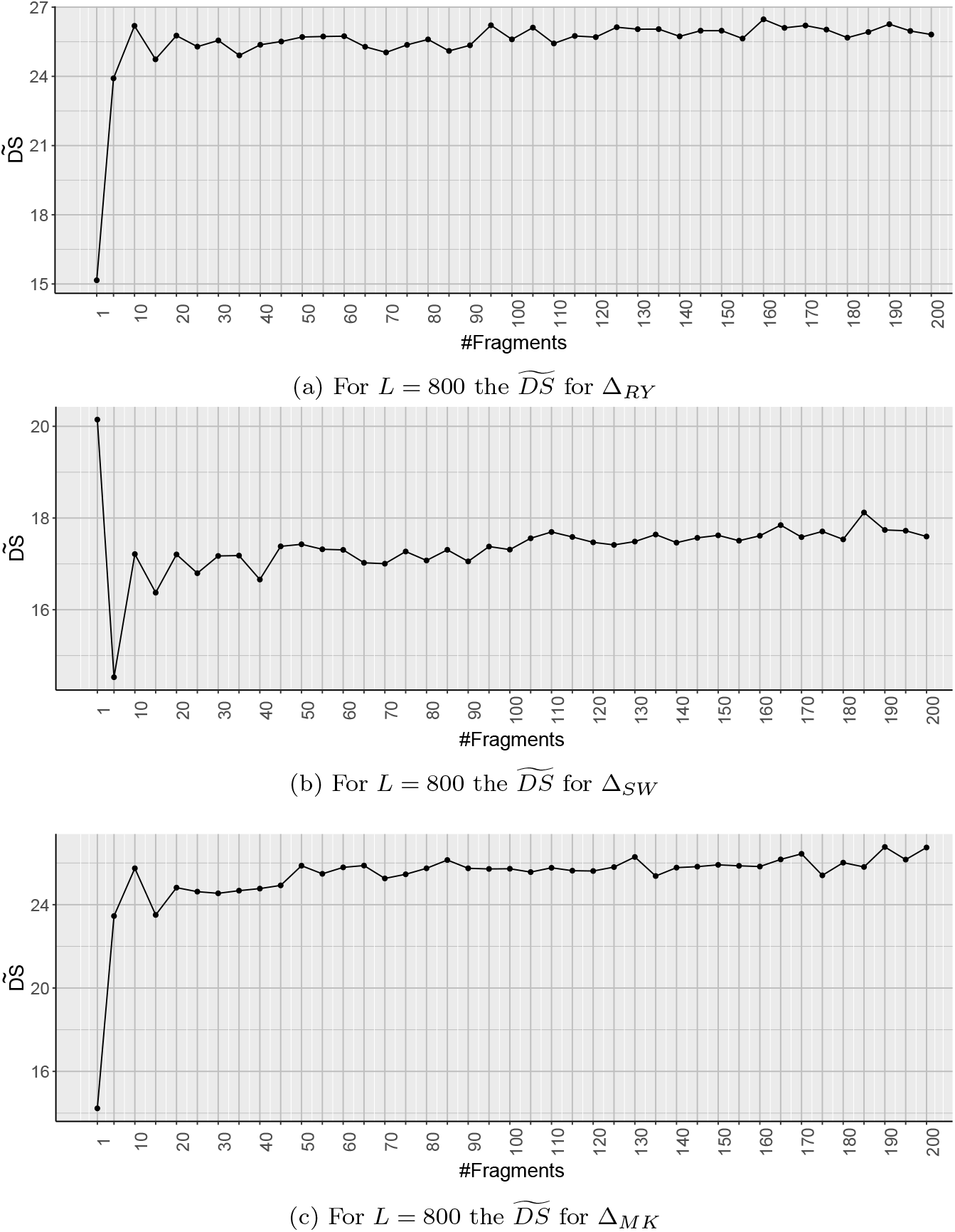
The *widetildeDS* for *L* = 800 for Δ_*RY*_, Δ_*SW*_, and Δ_*MK*_, respectively.

As the effect of *μ_intraclass_* is scaled down in 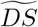, so 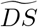 represents the effect of *μ_interclass_* on *DS* for the corresponding value of 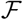. It is also shown in Fig. 4 (b, d, and f) that, for *L* = 800, after a period the change of 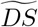 becomes less than 5%. We consider that as the stable state of 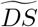. This implies that after a certain value of 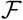, the interclass distance does not increase significantly with an increasing value of 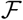. Within this *stable state*, there are some segments where the changes of 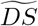 are less than 2%. We consider those as *stationary regions.* We consider the *DS* (obtained from Eq. (13)) of those stationary regions. We choose that value of 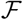 for which the maximum value of *DS* lies within these stationary regions. Empirically it is tested that for that value of 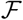, GRAFree infers tree with better clades for both GFP-RY, GFP-SW, and GFP-MK. Hence, it is considered that for the particular value of 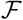 the feature vector of the species represents the Δ better for comparative genomic study. Hence we select the value of 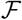 as 160, 165, and 165 for Δ_*RY*_, Δ_*SW*_, and Δ_*MK*_, respectively.

### Selection of the value of *α*

For a given *L* and 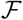, to choose the value of *α* within the range of [0,1], we consider the distance matrix and apply the same concept over the scaler values of the distance matrix to compute *DS* (refer to Eq. (13)). We derive the mean and variance of all pairwise distances, say, *μ_i_* and 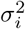 respectively, between the species of class *i*. Similarly, compute the *μ_intraclass_* as the mean of intraclass variances using the Eq. (10).

The mean of interclass distance is derived as following,

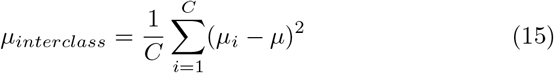

It is also be observed that the value of *DS* becomes stabilized after a period. Now we choose that value of *α* for which the maximum value of *DS* is obtained from Eq. (13) after stabilized. From Fig. 5, it can be observed that for all the selected values of 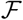 (−160, 165, and 165 for Δ_*RY*_, Δ_*SW*_, and Δ_*MK*_, respectively) and *L* (=800), the maximum value of *DS* is obtained for *α* =1.

**Figure 5:**
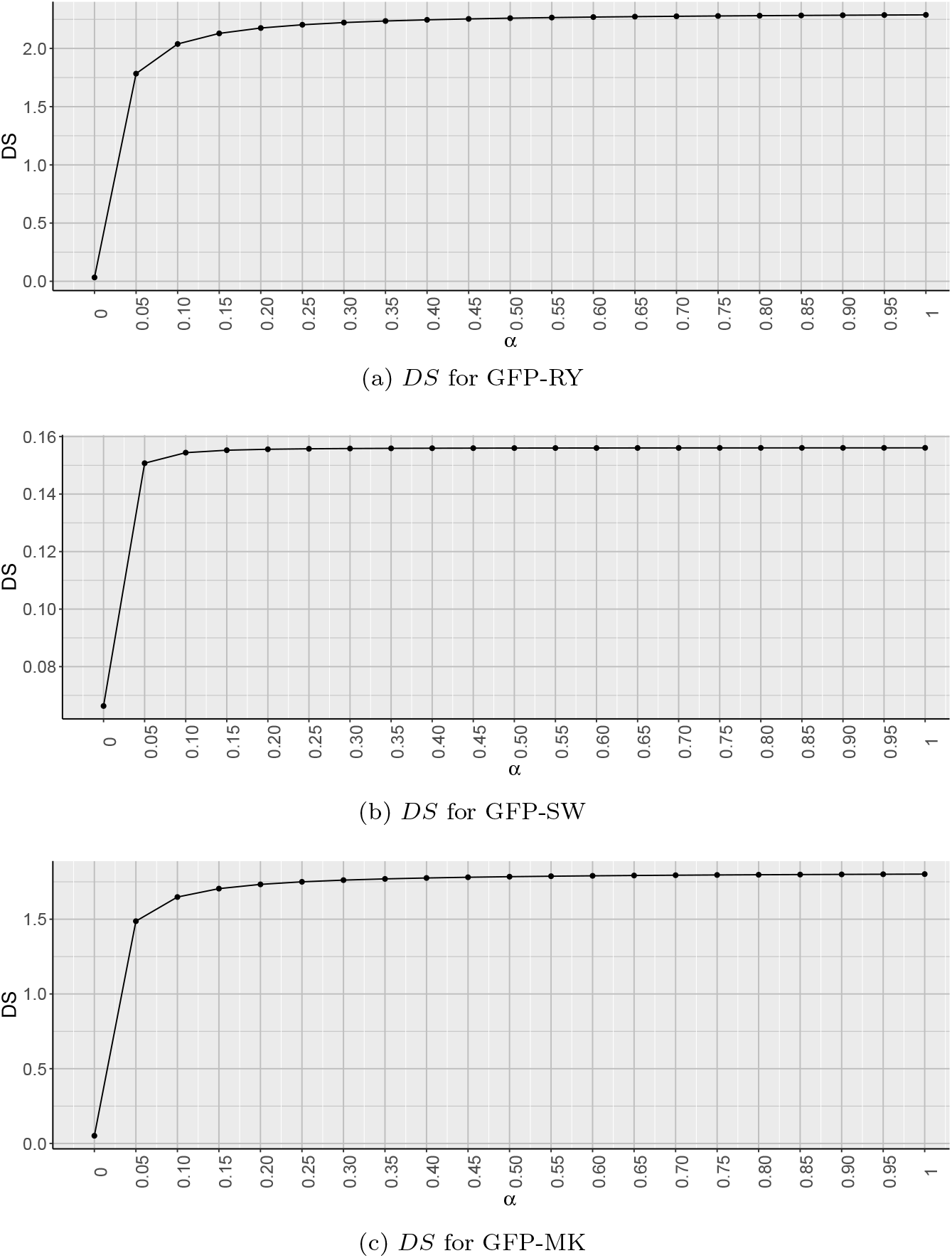
The *DS* for all the values of α from 0 to 1, given *L* − 800 and 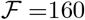, 165, and 165 for GFP-RY, GFP-SW, and GFP-MK, respectively.

Hence, we consider *L* = 800, 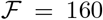, and *α* = 1 as the value of the parameters to derive the phylogenetic tree for GFP-RY. Similarly, for GFP-SW and GFP-MK, the value of *L*, 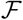, and *α* are chosen as 800, 165, 1 and 800, 165, 1, respectively. It is also noted empirically that the tree inferred using these parameters accumulates most of the intraclass species within same clade.

### Performance measure

In this study, we consider four different classes with more than 50 families of species. For measuring the accuracy of the derived tree we consider four classes, seven orders, and four families which are monophyletic and have more than ten representative species in our dataset. Our primary objective of this proposed method of performance measuring is to cluster the monophyletic species according to their respective class, order, or family. This is a quantitative measure of the deformation of a given monophyletic clade of phylogenetic tree. This measurement is useful especially when we do not have the reference tree to compare. The *transfer level (TransLv)* proposed here is defined as the minimum number of levels require to move a species to another target clade. The objective behind the transfer of a species to another clade is either to place the species to its appropriate clade or to remove the species from an inaccurate clade. The *total transfer level (TTL)* of a clade is the sum of the transfer levels to make a clade correct. The proposed measure of accuracy of a clade is based on the total transfer level of the clade. Using the TTL we compute the proposed measure, *Deformity Index*, which is a quantitative measure of the deformation the clades of the tree. Hence, for the ideal case where each species is placed within the proper clade, the value of Deformity Index is zero. The computation of the Deformity Index of a clade is described in Algorithm 1.

## Results and discussion

### Bootstrapping

The conventional method of bootstrapping [17], [14] considers the aligned sequences to resample and replicate. As we are developing an alignment-free method of phylogeny construction, the conventional bootstrapping method may not be applicable for this case. The main motivation of bootstrapping is to generate the population from a single genome. It is observed that the average intraspecific genetic variation is within 1% [38], [70]. So here we propose a bootstrapping technique which considers the genetic variance of a sample space within 1%. For that we apply mutations at each location with a probability of 1% and consider an unbiased selection of the nucleotides at each location. We generate 100 replicas using this bootstrapping method and construct trees from each set of sequences using GRAFree method by setting values of *L*, 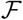, and *α* as discussed in the previous section. Felsenstein’s bootstrapping method [17] assesses the robustness of phylogenetic trees using the presence and absence of clades. For the large scale genomics Felsenstein’s bootstrap is not efficient to sum up the replicas. For the hundreds of species this method is inclined to produce low bootstrap support [36]. So here we apply a modification of Felsenstein’s bootstrapping, where the presence of a clade is quantified using the transfer distance proposed in [36]. The transfer distance [10] or R-distance [13] is the minimum number of changes required to transform one partition to other. We computed the occurrence of each clade using the tool BOOSTER [36].

**Algorithm 1.**
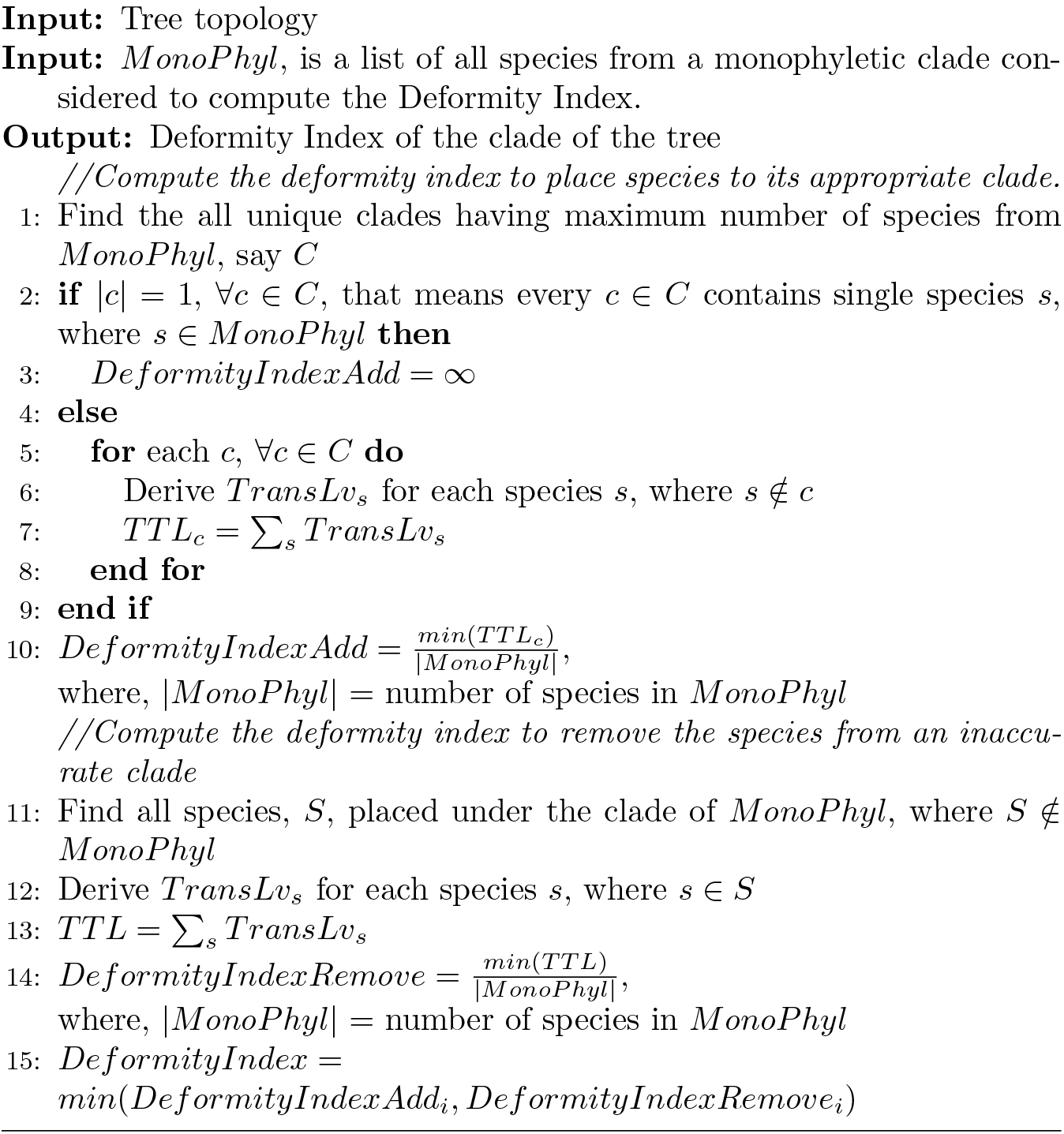
Algorithm for measuring the Deformity Index

### Observations from derived phylogenetic trees

Here, we present phylogenetic trees generated by our proposed method, GRAFree, using the whole mitochondrial genome sequences of the selected species. We consider the value of *L*, 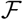, and *α* derived from the selection technique proposed in the previous section. It is observed in Table 2 that the average Deformity Index of GFP-RY (please refer to Eq. (1)) is lower than that of GFP-SW and GFP-MK (please refer to Eq. (2) and (3), respectively). These results infer that the skew (AG skew and CT skew) represented by the Eq. (1) bears the signature of genomic contents related to the evolution more precisely than that of the other skews. Hence, the skew of the genomic signature may also require careful investigation on the matter of evolutionary relationships. The 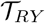, 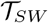, and 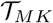 are shown in Fig. 6. Fig. 7 presents the final tree after merging all three cases using the COSPED-II algorithm.

**Table 2:** Deformity index for different trees

| Method | Class |  |  |  |
| --- | --- | --- | --- | --- |
|  | Mammalia | Avian | Fish | Insecta |
| GFP-RY | 1.87 | 0.39 | 0.25 | 3.92 |
| GFP-SW | 2.30 | 4.54 | 0.56 | 0.22 |
| GFP-MK | 1.57 | 4.54 | 6.50 | 7.75 |
| COSPED | 0.50 | 4.56 | 0.41 | 2.61 |
| **Reference Methods** |  |  |  |  |
| FFP | 11.75 | 12.86 | 12.59 | 102.25 |
| Chebyshev | 8.88 | 24.76 | 14.00 | 7.94 |
| Canberra | 0.00 | 0.00 | 0.31 | 0.00 |
| $D_2^*$ | 0.00 | 0.00 | 0.00 | 0.00 |
| Co-phylog | 0.00 | 0.00 | 0.00 | 0.00 |

| Method | Order |  |  |  |  |  |  |
| --- | --- | --- | --- | --- | --- | --- | --- |
|  | Carnivora<br>(Order of<br>Mammal) | Charadriiformes<br>(Order of<br>Bird) | Columbiformes<br>(Order of<br>Bird) | Gadiformes<br>(Order of<br>Fish) | Perciformes<br>(Order of<br>Fish) | Hymenoptera<br>(Order of<br>Insect) | Blattodea<br>(Order of<br>Insect) |
| GFP-RY | 1.79 | 6.89 | 5.96 | 1.16 | 0.25 | 3.43 | 3.63 |
| GFP-SW | 2.10 | 5.58 | 5.96 | 4.16 | 0.42 | 1.86 | 4.00 |
| GFP-MK | 1.41 | 9.42 | 5.67 | 2.89 | 0.17 | 4.29 | 3.00 |
| COSPED | 0.38 | 5.47 | 7.04 | 1.68 | 0.33 | 2.93 | 3.19 |
| **Reference Methods** |  |  |  |  |  |  |  |
| FFP | 11.63 | 132.00 | 10.14 | 12.11 | 74.67 | 295.64 | 201.38 |
| Chebyshev | 7.47 | 13.80 | 16.43 | 12.16 | 3.83 | 8.79 | 5.63 |
| Canberra | 0.13 | 1.75 | 0.00 | 0.47 | 0.00 | 0.00 | 3.13 |
| $D_2^*$ | 0.20 | 0.00 | 0.00 | 0.11 | 0.00 | 6.21 | 2.50 |
| Co-phylog | 0.07 | 1.75 | 0.00 | 0.11 | 0.00 | 0.00 | 0.31 |

| Method | Family |  |  |  |
| --- | --- | --- | --- | --- |
|  | Felidae<br>(Family of<br>Carnivora) | Threskiornithidae<br>(Family of<br>Pelecaniformes) | Gadidae<br>(Family of<br>Gadiformes) | Sciaenidae<br>(Family of<br>Perciformes) |
| GFP-RY | 4.21 | 43.50 | 0.40 | 0.73 |
| GFP-SW | 3.43 | 3.75 | 1.00 | 2.09 |
| GFP-MK | 3.29 | 3.75 | 1.70 | 0.45 |
| COSPED | 3.43 | 3.50 | 0.40 | 1.91 |
| **Reference Methods** |  |  |  |  |
| FFP | 10.14 | 276.50 | 11.00 | 82.45 |
| Chebyshev | 3.14 | 649.00 | 4.30 | 4.09 |
| Canberra | 0.29 | 0.00 | 0.00 | 0.00 |
| $D_2^*$ | 0.86 | 0.00 | 0.00 | 0.00 |
| Co-phylog | 0.29 | 0.00 | 0.00 | 0.00 |

**Figure 6:**
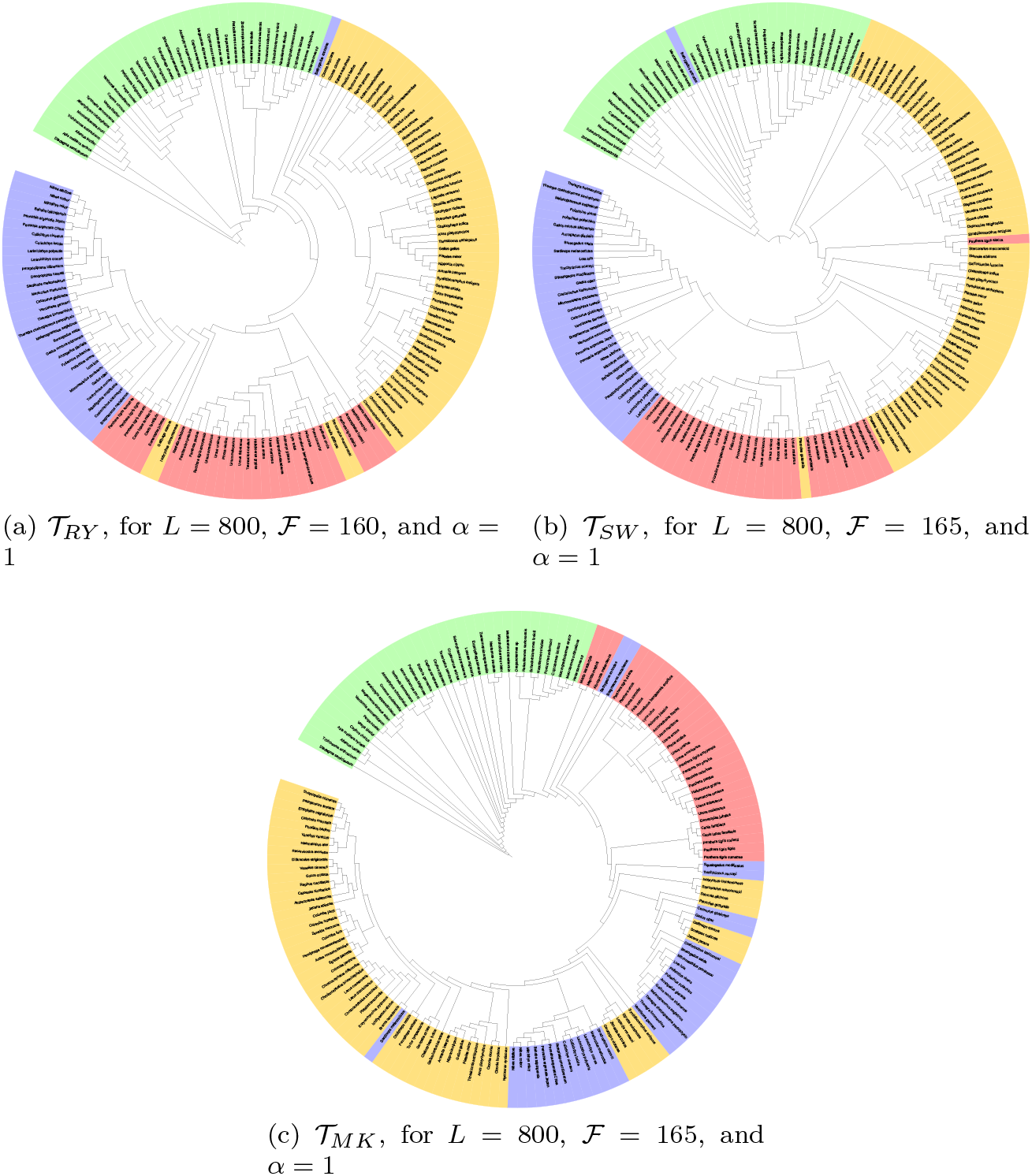
The inferred trees with the selected value of *L*, 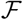, and *α* for GFP-RY, GFP-SW, and GFP-MK. The mammals, fishes birds, and insects are represented by red, blue, yellow, and green, respectively.

**Figure 7:**
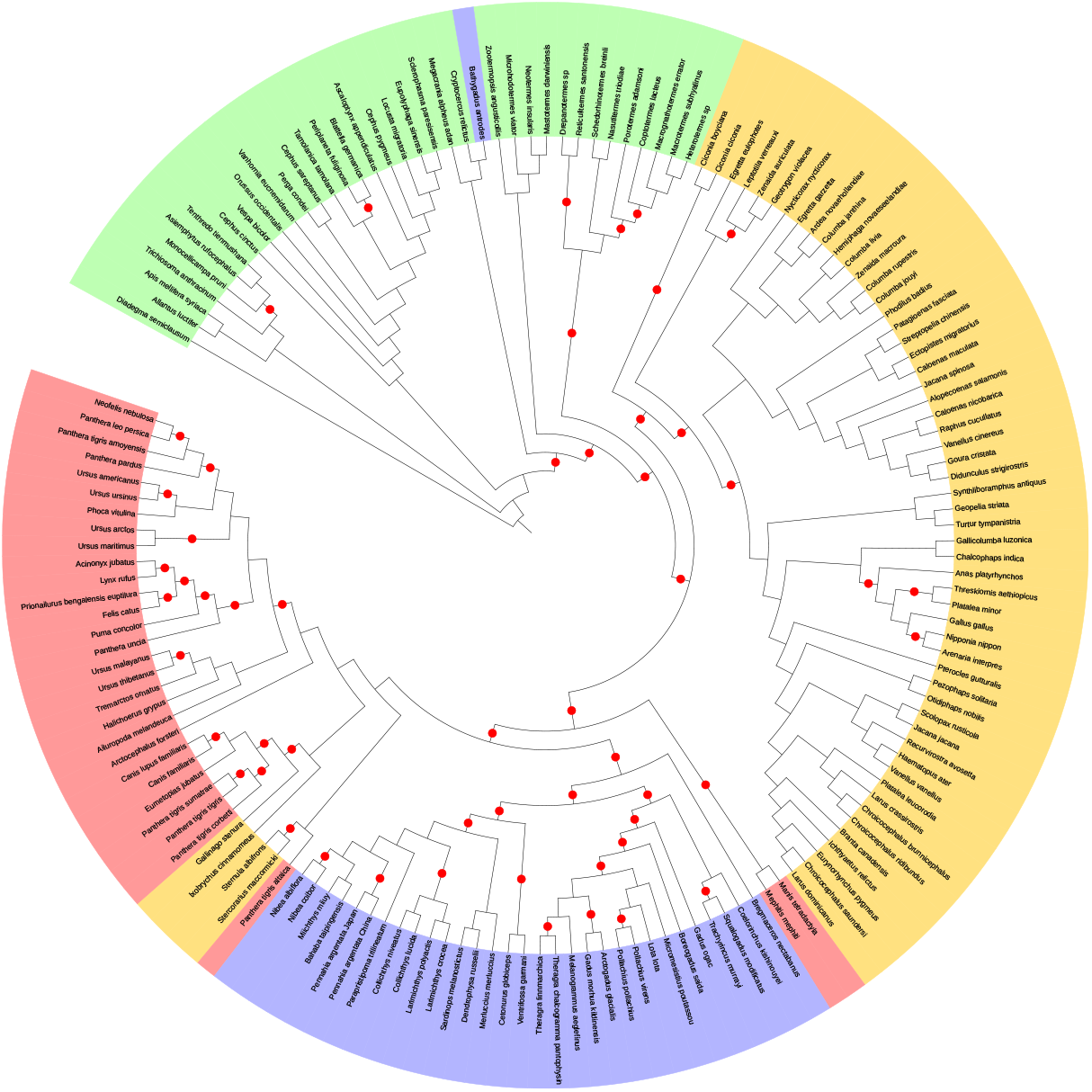
Inferred tree, 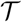 after merging 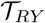, 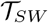, and 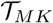. Red colored group represents the class mammals, blue colored group represents the class ray-finned fishes, yellow colored group represents the class birds, and green color represents the species from insect. The red dot on the branch represents the bootstrap score of that clade is greater than 75%.

To measure the performance of 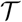, we chose 15 monophyletic clades of four classes, seven orders, and four families. It is observed that 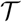 has formed the monophyletic clades for the three major classes, mammals, fishes, and birds with minor deviations, whether insects are inferred as paraphylic. This tree also infers insects as the oldest class among these four classes followed by birds, mammals, and fishes. Mammals and fishes are inferred as the sister clades in 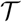. The deformity index of different trees are shown in Table 2.

### Observations from reference methods

We have examined five different existing distance measures of alignment free method for phylogenetic reconstruction, i.e. FFP [62], [63], 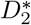 [59], [68], Chebyshev, Canberra, and Co-phylog [72] using the tool ACcelerated Alignment-FrEe sequence analysis (CAFE) [44] on our dataset. We measured each distances with the word (k-mer) length of 10. The resultant trees are shown in Fig. 8. It is found that 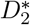 and Co-phylog separate four major clades accurately (with the average deformity index of 0), Canberra separate four major clades with minor errors (with the average deformity index of 0.11) in their inferred trees whereas the other methods, FFP and Chebyshev, cannot identify the four classes as clades. The bootstrap support of the clades of the inferred tree from 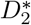, Co-phylog, and Canberra methods are also very high. The 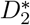 and Co-phylog infer the tree having insects as the oldest clade followed by fishes, birds, and mammals. Mammals and birds are inferred as the sister clades in the derived trees from both 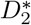 and Co-phylog methods. So the it is accepted that the 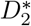, Co-phylog, and Canberra infer better clades than that of FFP and Chebyshev.

**Figure 8:**
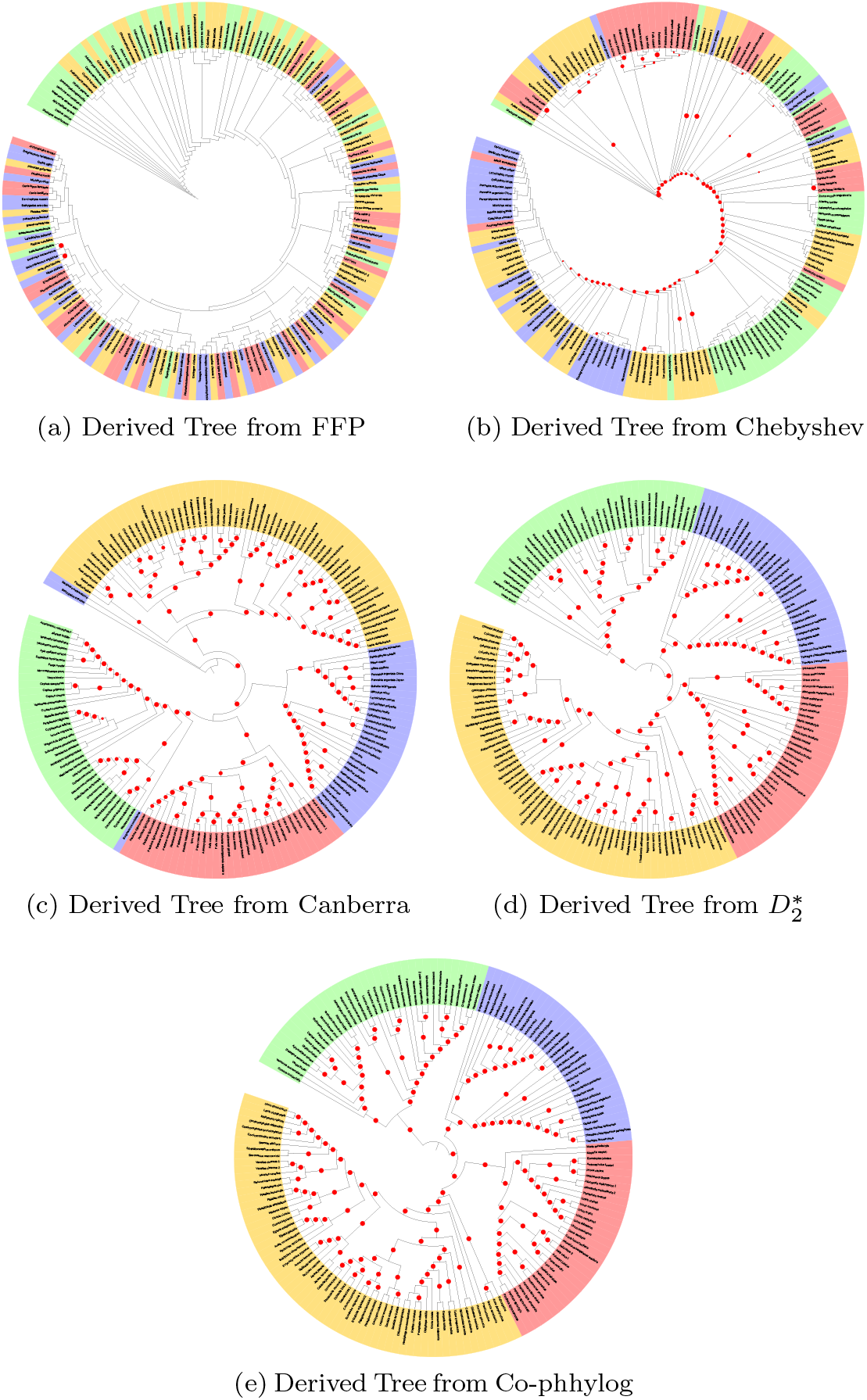
Derived tree from the reference methods. The mammals, fishes birds, and insects are represented by red, blue, yellow, and green, respectively. The clades having the bootstrap scores more than 75% are denoted by the red dots on the branch.

### Complexity analysis

To compute the complexity of GRAFree, we consider *M* as the length of the genome sequences of two species, 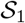 and 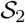, the length of the window to compute drift is *L*, and the number fragments of the drift is 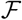. The GRAFree consists of three major steps.

#### Computing drift

As drift is computed considering two point coordinates on the GFP, so for each species the time complexity to compute drift is 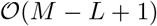. The drift sequence contains the 2D coordinate of total *M* − *L* + 1 points. So the space complexity of drift sequence of a species should be 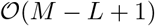.

#### Computing feature vector

Each fragment of the drift is represented by *μ*, Λ, λ, and *θ* (please refer to Subsection 1). The time complexity of Λ and λ are depending on the covariance matrix of drift. Since, we consider the 2D coordinate points in drift, the time complexity of computing Λ and λ for each species is 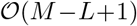. Time complexity of computing *μ* and *θ* are linearly related to the length of drift sequence. Hence, for each species the time complexity of computing the feature vector for 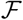 number of fragments is 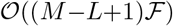. Similarly, the space complexity of feature vector for each species is 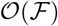.

#### Computing distance between a pair of species

GRAFree considers the distance function which computes the distances for all 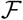 number of fragments in a constant time, means 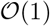. So the time complexity of computing distance between a pair of species is 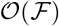.

Hence, both time and space complexity of GRAFree is 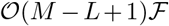.

#### Complexity analysis of FFP

FFP considers the frequencies of all k-mers. For the *M* length of sequence, the time complexity to compute the feature frequency of all k-mers is 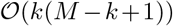. FFP uses Jensen-Shannon Divergence (JSD) for computing the distance between two feature frequency profiles. As JSD consider the entropy for deriving the distance, hence for (*M* − *k* + 1) length of sequences the time complexity for computing the JSD is 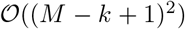. So the total time complexity of FFP for computing distance between two sequences is 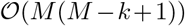. Since, FFP considers all k-mers to compute the feature frequency profile of the sequence, the total space complexity for nucleotide is not more than 4^*k*^. Hence, the space complexity of FFP for two sequences is 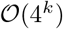.

#### Complexity analysis of Chebyshev

Chebyshev distance function considers the number of occurrences k-mers in the sequences. So for *M* length of sequences, the time complexity of computing the occurrence of a particular k-mers is 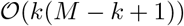. The maximum among the absolute value of the difference between each k-mers of two sequences is considered as the Chebyshev distance. So the time complexity of Chebyshev for computing the distance between two sequences is 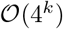. Since, Chebyshev considers the occurrence of all k-mers, the space complexity of Chebyshev for two sequences is 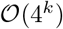.

#### Complexity analysis of Canberra

Similar to the Chebyshev, Canberra distance function also considers the number of occurrences of k-mers in the sequences. Hence, the time complexity to compute the occurrence k-mers is 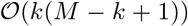. Canberra distance is considered as the summation of the ratio between the absolute value of the difference between each k-mer and the total occurrence of that particular k-mer within two sequences. Hence, the time complexity for computing the Canberra distance between two sequences is 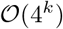. Similarly the space complexity of Canberra for two sequences is 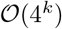.

#### Complexity analysis of 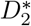

Similar to the FFP, 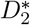 considers occurrence of k-mers in the sequences. So for *M* length of sequences, the time complexity of computing the occurrence of a particular k-mers is 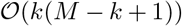. 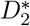 takes the probability of the k-mers within the combined sequence of the two sequences. For combined sequence of 2*M* length, the time complexity of computing the probability of each k-mer is 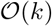. Using these two values, 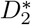 computes the distance for the particular k-mer. Finally, the sum of the distances for all k-mers is considered as the distance between two sequences. Hence, the time complexity of 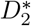 for computing the distance between two sequences is 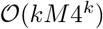. As it stores the occurrence of all k-mers, the space complexity is 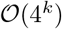.

For the large scale genomic study the time and space complexities are one of the important things to be remembered. We compare the time and space complexity of GRAFree with some of the existing methods in Table 3. It can be observed that GRAFree is significantly efficient than all the selected existing methods in order of the time and space complexity. It is noted that 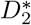 and Co-phylog are efficient in quality of reconstruction of tree. The execution time of different methods are shown in Fig. 9.

**Table 3:**
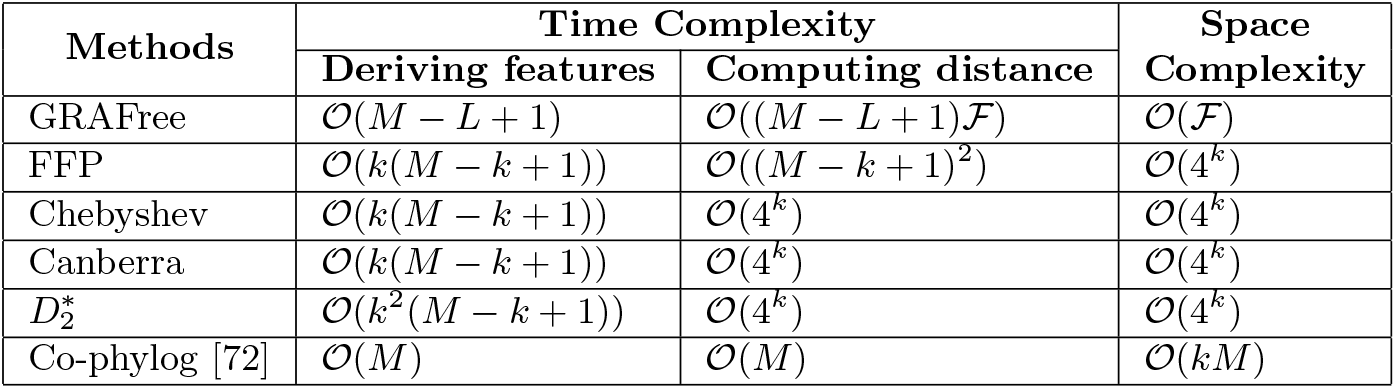
Time and space complexity to compute distance between two sequences by different methods

**Figure 9:**
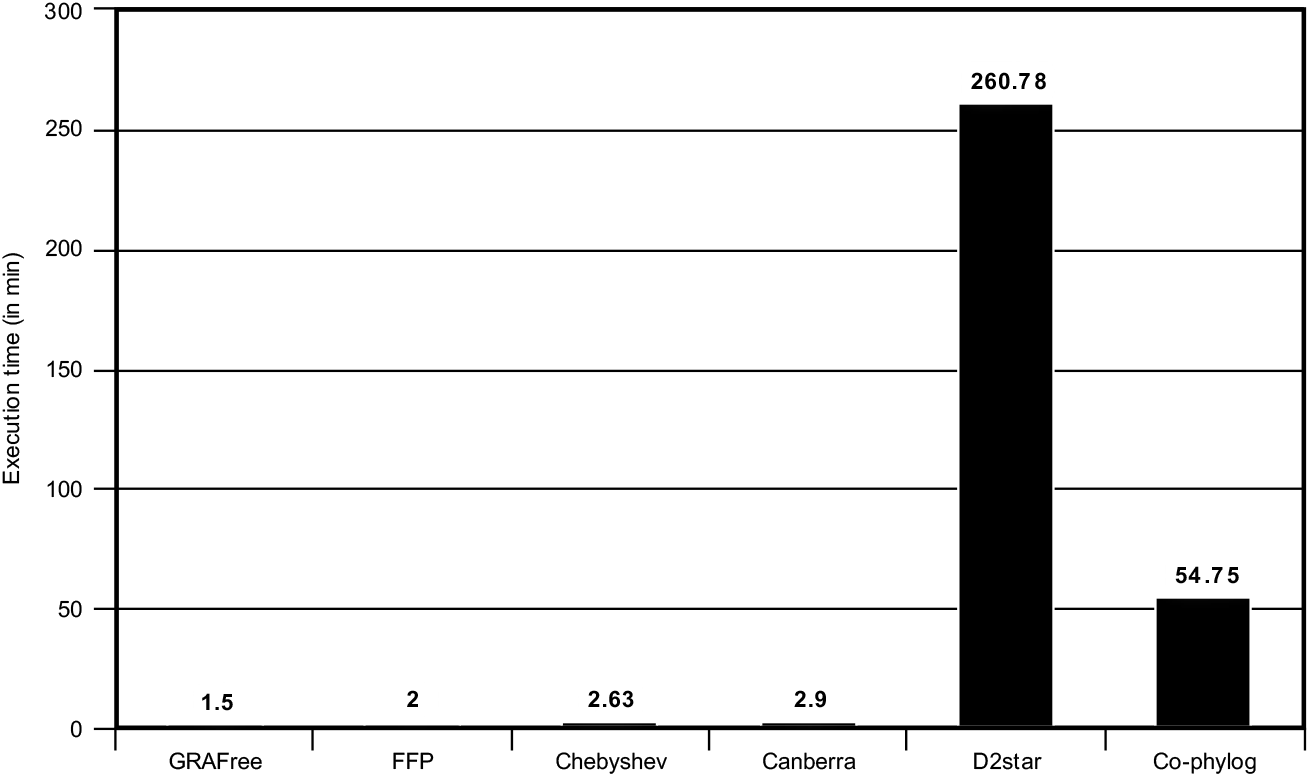
The execution time for different methods. All the methods are executed in the same system. The configuration of the system is 16GB RAM, Intel Core i5 processor, and it had 64 bit Ubuntu 17.10

## Conclusion

We have proposed a 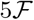-dimensional feature space and a new metric for capturing evolutionary relationship using large scale genomic features in the method GRAFree. GRAFree uses the graphical representation of the genome. In this study we have selected three very naive graphical representations of a genome considering residues independently. We have also proposed a novel measure to evaluate the performance of the techniques. The resultant tree accumulates most of the monophyletic clades with minor deviations. In spite of these limitations, we could observe presence of evolutionary traits in the proposed feature descriptor extracted from the whole mitochondrial sequences. The tree has a high bootstrap support for a good number of clades. These demonstrate the effectiveness of the proposed feature representation, as well as the metric for measuring the pairwise distances of species.

## Footnotes

1 Website of NCBI database: http://www.ncbi.nlm.nih.gov

2 Website of “NCBI Help Manual”: http://www.ncbi.nlm.nih.gov/books/NBK3831/

